# Effect of genomic and cellular environments on gene expression noise

**DOI:** 10.1101/2022.08.31.506082

**Authors:** Clarice KY Hong, Avinash Ramu, Siqi Zhao, Barak A Cohen

## Abstract

Individual cells from isogenic populations often display large cell-to-cell differences in gene expression. This “noise” in expression derives from several sources, including the genomic and cellular environment in which a gene resides. Large-scale maps of genomic environments have revealed the effects of epigenetic modifications and transcription factor occupancy on mean expression levels, but leveraging such maps to explain expression noise will require new methods to assay how expression noise changes at locations across the genome. To address this gap, we present Single-cell Analysis of Reporter Gene Expression Noise and Transcriptome (SARGENT), a method that simultaneously measures the noisiness of reporter genes integrated throughout the genome and the global mRNA profiles of individual reporter-gene-containing cells. Using SARGENT, we performed the first comprehensive genome-wide survey of how genomic locations impact gene expression noise. We found that the mean and noise of expression correlate with different histone modifications. We quantified the intrinsic and extrinsic components of reporter gene noise and, using the associated mRNA profiles, assigned the extrinsic component to differences between the CD24^+^ “stem-like” sub-state and the more “differentiated” sub-state. SARGENT also reveals the effects of transgene integrations on endogenous gene expression, which will help guide the search for “safe-harbor” loci. Taken together, we show that SARGENT is a powerful tool to measure both the mean and noise of gene expression at locations across the genome, and that the data generated by SARGENT reveals important insights into the regulation of gene expression noise genome-wide.

## Introduction

Gene expression is noisy, even among individual cells from an isogenic population^1^. Noisy gene expression leads to variable cellular outcomes in differentiation^2–5^, the response to environmental stimuli^6,7^, viral latency^8^, and chemotherapeutic drug resistance^9–11^. Explaining the causes of noisy expression remains an important challenge.

A gene’s genomic environment, defined here as the composition of nearby *cis*-regulatory elements and local epigenetic marks, can influence its expression noise^12^. Some features of genomic environments that can affect noise include enhancers, histone modifications, and transcription factor (TF) occupancy^13–19^. These observations raise the possibility that genome-wide patterns of expression noise could be explained using the large-scale epigenetic maps that have proved useful in explaining mean expression levels^20–22^. Leveraging these resources to explain expression noise will require maps of the genome that show the influence of diverse genomic environments on this noise. Producing these maps will require new experimental approaches, because the existing studies demonstrating the effects of epigenetic marks on expression noise have either been performed on endogenous genes, where the effects of different chromosomal locations are confounded with the effects of the different endogenous promoters or rely on low-throughput imaging methods. Dar *et al*. assayed the noisiness of large numbers of genomic integrations but was unable to assign genomic locations to the measured reporter genes^16^. Two other studies have assayed integrations in a high-throughput manner but measured protein levels by flow cytometry rather than mRNA levels^23,24^. Thus, we still lack a high-throughput, systematic way of quantifying the impact of genomic environments on expression noise.

In addition to intrinsic features such as the local genomic environment, extrinsic features, such as the global cellular state of a cell, can also influence gene expression noise^25–29^. For example, variation in the cell cycle, cell size, or signaling pathways can all impact gene expression noise^1,30,31^. However, the relative contributions of intrinsic vs extrinsic features on gene expression noise in mammalian cells remains unclear.

Here we report Single-cell Analysis of Reporter Gene Expression Noise and Transcriptome (SARGENT), a highly parallel method to measure the mean and noise of a common reporter gene that has been integrated at locations across the genome. Analysis of SARGENT data showed that different histone modifications explain the mean and noise produced across the genome. In SARGENT, multiple reporters are integrated in each cell, allowing us to separate the intrinsic and extrinsic contributions to noise. Sequencing the associated single-cell mRNA transcriptomes further enabled us to attribute the extrinsic noise to differences in the cellular substates between isogenic cells. To our knowledge, this is the largest genome-wide survey of the impact of intrinsic and extrinsic noise in gene expression. Taken together, our results show that SARGENT is a powerful tool to study how genomic environments and cellular context control expression noise.

## Results

### A high-throughput method to measure mean and noise across the genome

We developed a high-throughput method to test the effects of genomic environments on the mean and noise of gene expression. Our goal was to integrate a common transgene across the genome and then, for individual cells, measure both the transcripts produced from the transgene and the global mRNA profile. This allows us to compute the mean and noise of reporter gene expression at each location, and correlate reporter gene expression with the cellular mRNA state of each cell. Because every unique integration contains the same transgene, the measured differences in the mean and noise of reporter gene expression are directly attributable to the influence of genomic environments or cellular states.

We first generated a reporter gene with a library of 16bp random barcodes (Location barcode, locBC) in its 3’UTR **(Figure 1)**. Due to the diversity of the locBCs, each locBC is only associated with a single location in the genome^21^. The reporter gene consists of a Cytomegalovirus (CMV) promoter driving the expression of a fluorescent protein and contains a capture sequence from the 10x Genomics Single Cell Gene Expression 3’ v3.1 with Feature Barcoding Kit. The 10x gel beads contain both the complementary capture sequence and polyT sequences, allowing us to isolate the transcripts produced from the reporter gene and the cellular transcriptome.

**Figure 1:**
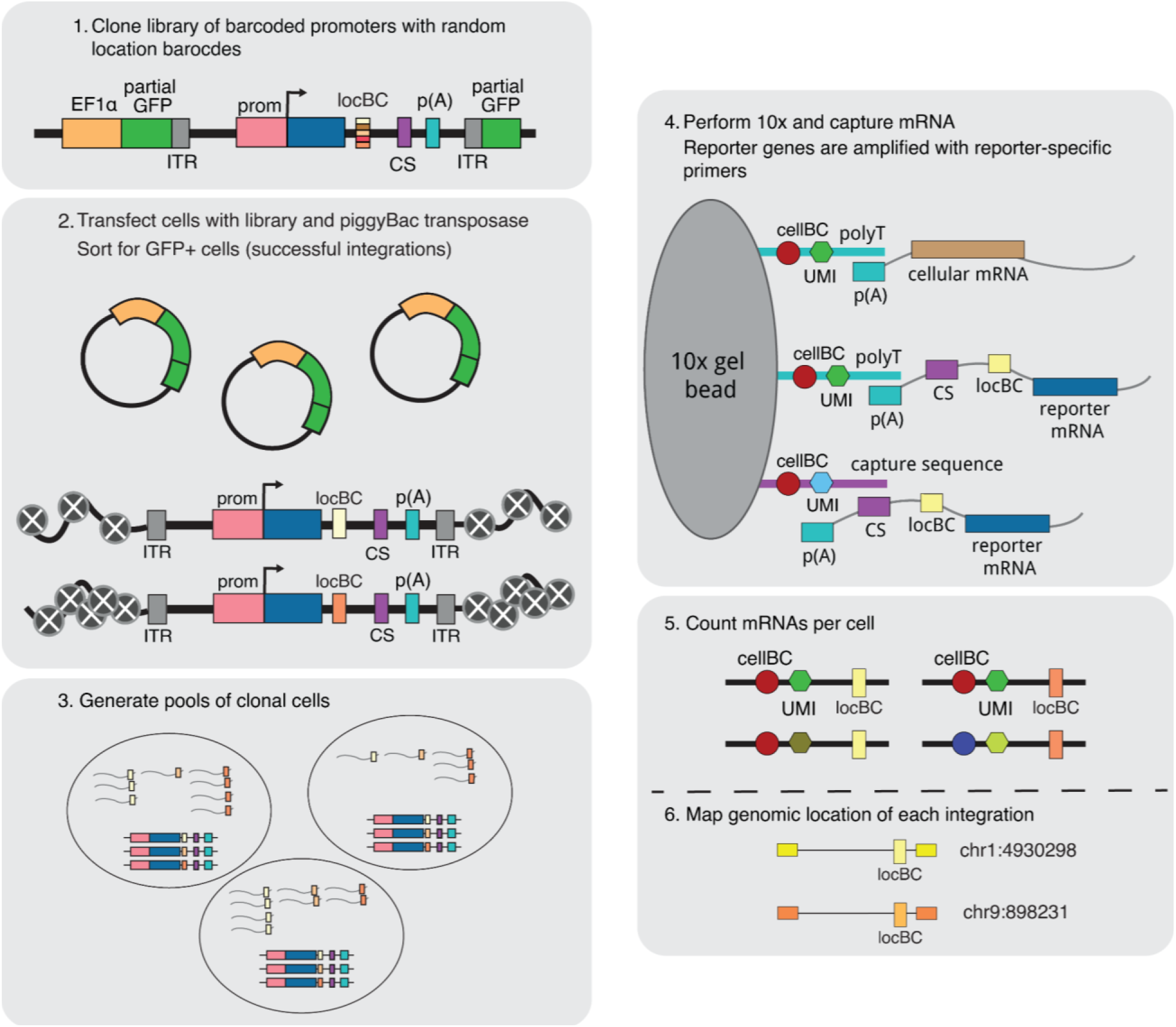
Overview of the SARGENT workflow. In step 1, a reporter gene driven by the CMV promoter is randomly barcoded with a diverse library of location barcodes (locBC) upstream of the 10x capture sequence (CS). The reporter genes are randomly integrated into K562 cells and sorted for cells with successful integrations (step 2), then sorted again after a week into pools to ensure that each barcode is only represented once per pool (step 3). We then performed scRNA-seq to capture the transcriptome and amplify the expressed barcodes from integrated reporter genes (step 4). The number of expressed barcodes per cell were then tabulated (step 5). To identify the genomic locations of the integrations we also mapped the location of each locBC in bulk (step 5). (ITR = Inverted terminal repeat; prom = promoter)

To generate chromosomal integrations across the genome we cloned the reporter gene library onto a piggyBac transposon vector. The library was transfected into cells along with piggyBac transposase to allow random integrations of the reporter into the genome. After selecting for integrations in K562 cells, we mapped the locations of each integrated reporter (IR) and assigned each locBC to a specific genomic location. We then captured the reporter gene transcripts from single cells and amplified the barcodes (10x cell barcode, UMI, and locBC) using primers specific to our reporter gene (**Methods**). After sequencing and tabulating the mRNA counts for each IR we computed the expression level of the reporter gene at each genomic location in each single cell. For a subset of cells we also sequenced the mRNA profiles to simultaneously reveal the cell state of each individual cell.

### SARGENT measurements are accurate and reproducible

We performed SARGENT in K562 cells because of the abundance of public epigenetic data available for this cell line. We first assessed the reproducibility of the SARGENT method. Because replicate infections result in pools of cells with insertions at different genomic locations, we could not assess the reproducibility of independently transfected pools of cells. Instead, we assessed the reproducibility of SARGENT by growing the same pool of insertions twice and performing the SARGENT workflow independently on each sample. We detected 589 identical IR locations in both replicates, which represented 96% of the total IRs observed in both replicates. After quality control, we obtained data from 7680 single cells across replicates, and a total of 2,940,912 unique molecular identifiers (UMIs) representing expressed barcodes from the IRs in these cells. The replicates were well correlated for measurements of both mean and noise measured at each IR location (**Figures 2A, B**, mean Pearson’s *r* = 0.76, noise Pearson’s *r* = 0.72) indicating that measurements obtained by SARGENT are reproducible.

**Figure 2:**
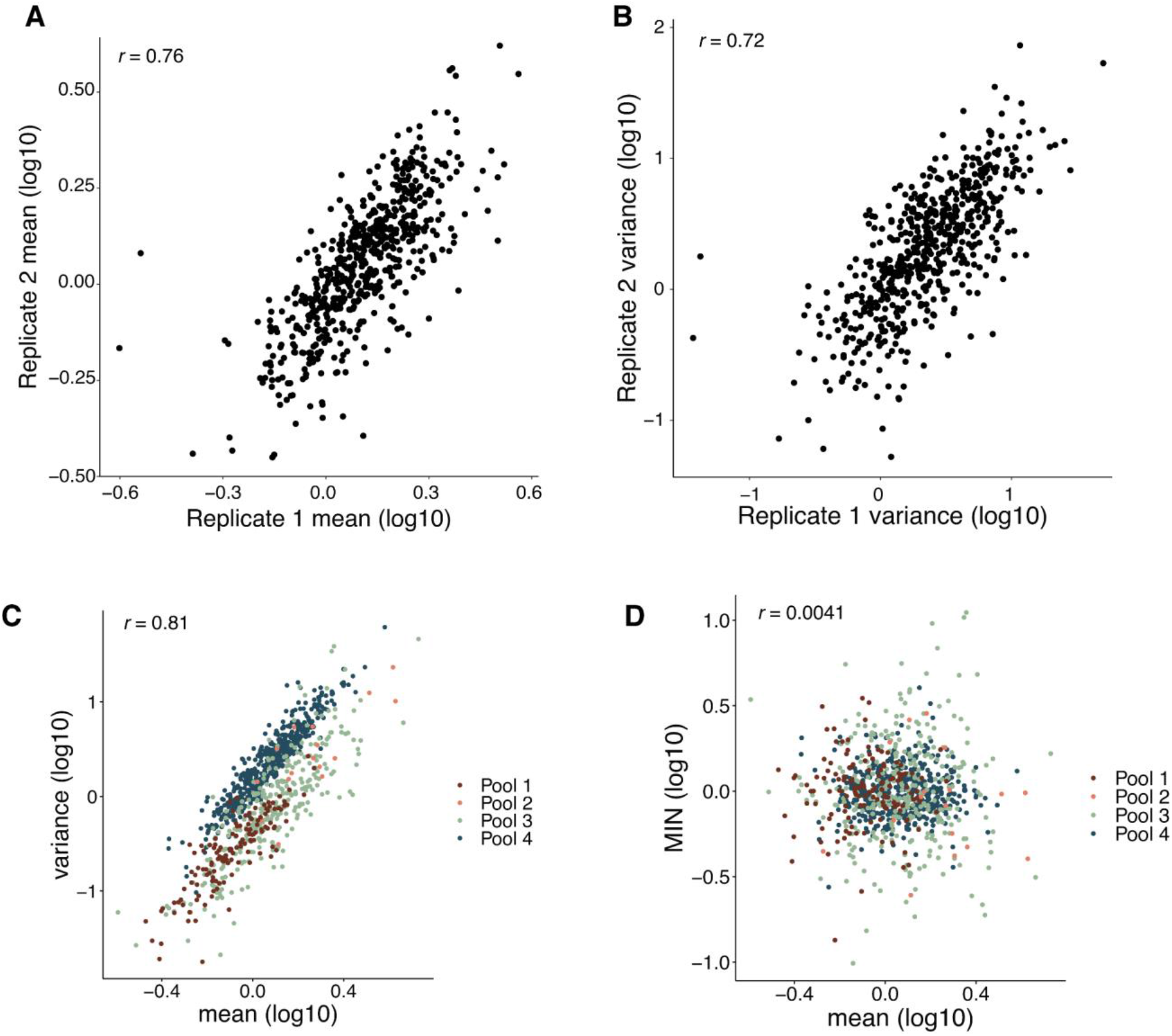
SARGENT measurements are accurate and reproducible. **(A)** Correlation of mean levels between technical replicates. **(B)** Correlation of variance measurements between replicates. **(C)** Mean and variance are correlated within each experiment. **(D)** Mean-independent noise corrects for mean effects on variance. Correlations shown are Pearson’s correlation coefficients (Pearson’s *r*).

To validate the single-cell measurements made by SARGENT, we also performed single-molecule Fluorescence *in situ* Hybridization (smFISH) on two known locations. The measurements of mean and noise made by smFISH agree with the SARGENT measurements for those locations **(Supplementary Figure 1)** demonstrating that our method is accurate and reproducible for measuring the mean and noise of expression.

### Measurements of mean-independent noise across different chromosomal environments

In total, we performed four experiments and generated mean and noise measurements for 939 integrations **(Supplementary Table 1)**. The integrations were spread across the genome and found in regions with different chromHMM annotations^32^ **(Supplementary Figures 2A, 2B)**, allowing us to study the effects of diverse chromosomal environments on expression noise.

The mean and noise of expression are often highly correlated^33,34^. Similarly, we found a strong correlation between the mean and noise in SARGENT data, indicating that a large proportion of an IR’s noise is explained by its mean level of expression **(Figure 2C)**. To identify chromosomal features that control expression noise independent of mean levels we regressed out the effect of mean levels on noise, leaving us with a metric we refer to as mean-independent noise (MIN)^33^. By design, MIN levels of IRs are uncorrelated with their mean expression levels **(Figure 2D)** whereas other measures of noise, such as the coefficient of variance or the Fano factor, retain residual correlation with mean levels in our data **(Supplementary Figures 2C, 2D)**. Thus, we used MIN as a measure of expression noise for all following analyses.

### Expression mean and noise are associated with different chromosomal features

We sought to identify chromatin features that would explain differences in MIN levels between genomic locations. Studies of genome-wide chromatin features in many cell lines and tissues have shown that the mean expression of a gene is correlated with its surrounding chromatin marks^21,35^. Thus, we asked whether chromatin features might also explain patterns of MIN across the genome. We split the IRs into bins of high or low mean levels, or high or low MIN levels, and identified chromatin features that were enriched in specific bins. As expected, IRs with high mean expression had higher levels of active chromatin marks such as H3K27ac, H3K4 methylation, H3K79me2 and H3K9ac **(Figure 3A)**. Conversely, IRs with high MIN did not exhibit significant differences between H3K27ac or H3K4me1 levels, and low MIN locations showed slightly elevated levels of H3K4me2/3, H3K79me2 and H3K9ac **(Figure 3B)**. These results suggest that different chromatin modifications influence the mean and noisiness of expression, and that more active genomic locations might also reduce MIN. This observation is consistent with previous studies showing that repressed chromatin is associated with high MIN^19,23^.

**Figure 3:**
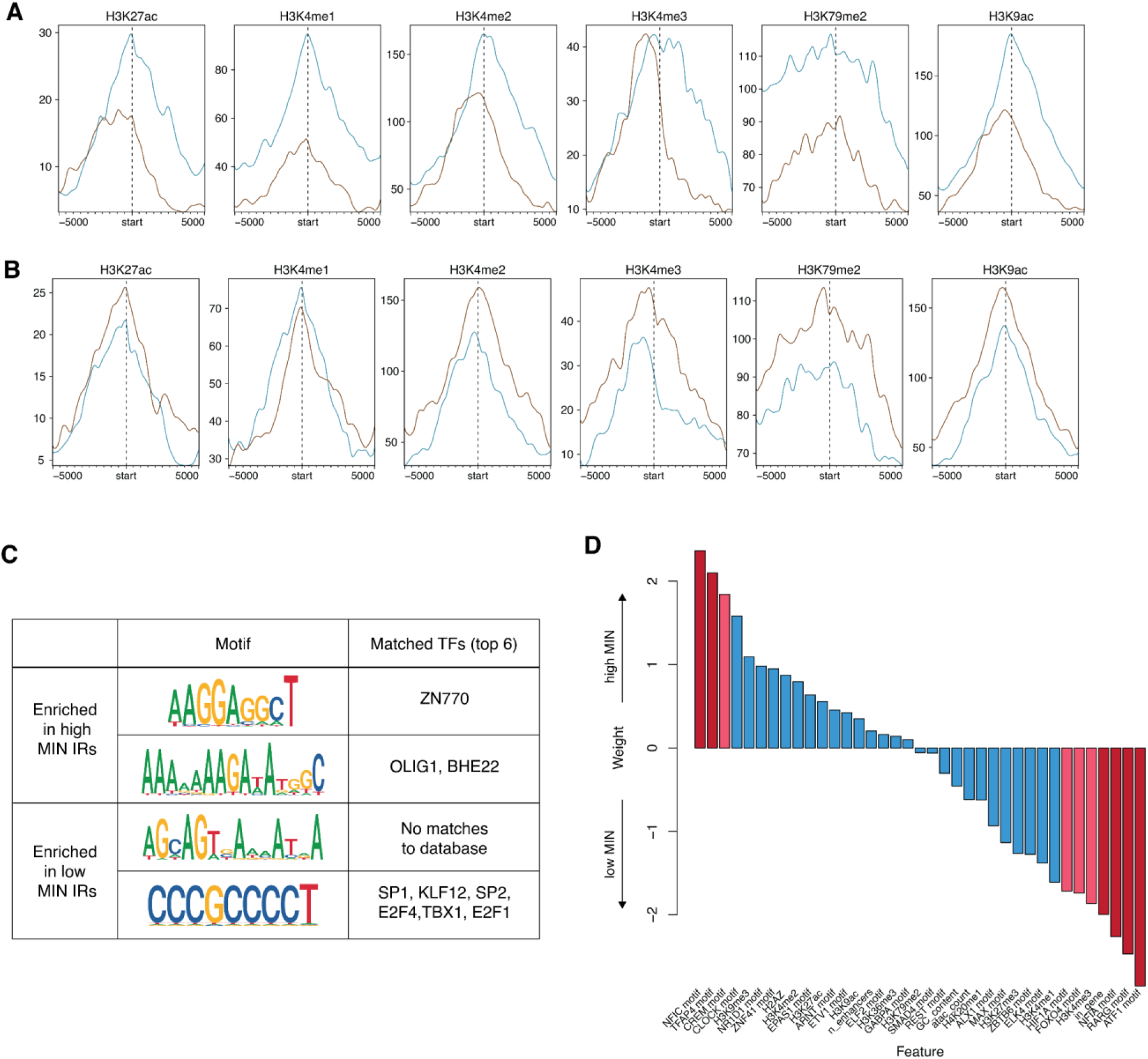
Expression mean and noise are associated with different chromosomal features. (A) Active histone modifications associated with high or low mean IRs. Start indicates the location of the IR, and each location was extended 5kb on either side. (B) Active histone modifications associated with high or low MIN IRs are different from those associated with mean. (C) Motifs enriched in high or low MIN IRs respectively, and potential TFs that match these discovered motifs. (D) Logistic regression weights of various intrinsic features associated with high or low MIN IRs. Red bars: p-value < 0.05; Pink bars: 0.05 < p-value < 0.1 from the logistic regression model.

The binding of TFs also impacts noise in gene expression. To identify TFs that might affect noise, we identified TFs whose occupancy is enriched near either high or low MIN IRs. Sequences at low MIN IRs are enriched for transcriptional activators such as SP1 and E2F4, while sequences at high MIN IRs are enriched for other TFs including TFs containing basic helix-loop-helix (bHLH) domains **(Figure 3C)**, suggesting that the cofactors recruited by different TFs have separable effects on expression mean and noise.

To assess the power of genomic features to predict the MIN of IR locations we trained a logistic regression model using chromatin modifications and DNA sequence features to classify high and low MIN locations and achieved an accuracy of 76%. When applied to data from a different pool, the trained model achieved 67% accuracy. The features with significant weights are the H3K4me3 mark, TF motifs (RARG, FOXO4, HIF1A, TFAP4, CREM, ATF1, NFIC, and NFIA) and whether the IR location was inside a gene (**Figure 3D, Supplementary Table 2**). Being inside a gene reduced the probability of being a high noise lR location, which could be due to local regulatory elements that dampen the noisiness of a gene’s expression. Similar to our results above, lower H3K4me3 increased the probability of being a high noise IR location. H3K4me3 is associated with active chromatin and supports the hypothesis that higher activity reduces IR MIN. Our observation is consistent with a previous study showing that H3K4me3 correlates with reduced noise at endogenous genes^19^. With respect to the effects of TFs on noise, the presence of some TF motifs increase the probability of being a high noise IR location (NFIC, CREM, TFAP4, CLOCK), whereas other TFs reduce the probability of being a high noise location (RARG, NFIA, ATF1, FOXO4, HIF1A).

We used a similar logistic regression framework to identify features that separate IR locations with high or low mean levels of expression and achieved an accuracy of 82%. When applied to holdout data, the trained model achieved 62% accuracy. The chromatin features that increase the probability of being a high mean IR location are lower levels of H3K27me3, lower levels of H3K4me2 and a higher number of ATAC-seq peaks, which agrees with the known effects of these features in bulk mean expression. The motifs that increased the probability of being a high mean IR location are higher numbers of motifs of the ZNF76, BACH1 and E2F3 TFs and fewer instances of the E2F7, SMAD3 and SOX5 motifs. (**Supplementary Figure 3, Supplementary Table 3**). Comparisons of the models explaining either mean or noise again show that different genomic features are correlated with gene expression mean and noise.

### Intrinsic and extrinsic factors have similar effects on gene expression noise

Expression noise caused by fluctuations in global factors affects all genes and is referred to as extrinsic noise, whereas intrinsic sources of noise are specific to individual genes^23,28–33^. The correlation between identical reporter genes in the same cell measures the balance between extrinsic and intrinsic noise, with extrinsic factors increasing the correlation^25^. In SARGENT, the correlation between IRs in the same cells is a measure of extrinsic factors that affect noise across IR locations.

For our analysis of extrinsic noise we first identified IRs in the same clonal cells using the co-occurence of locBCs between single cells. We identified 192 clones, with a mean of three integrations per clone (**Supplementary Figure 4A, Supplementary Table 4**). Of these 192 clones, 45 contain more than one integration (**Figure 4B**), making them suitable for an analysis of extrinsic noise. To validate the identified clones, we individually mapped IR barcodes in sixteen clones and found that 94% of the individually mapped IR locations could be uniquely assigned to an identified clone **(Figure 4B)**.

**Figure 4:**
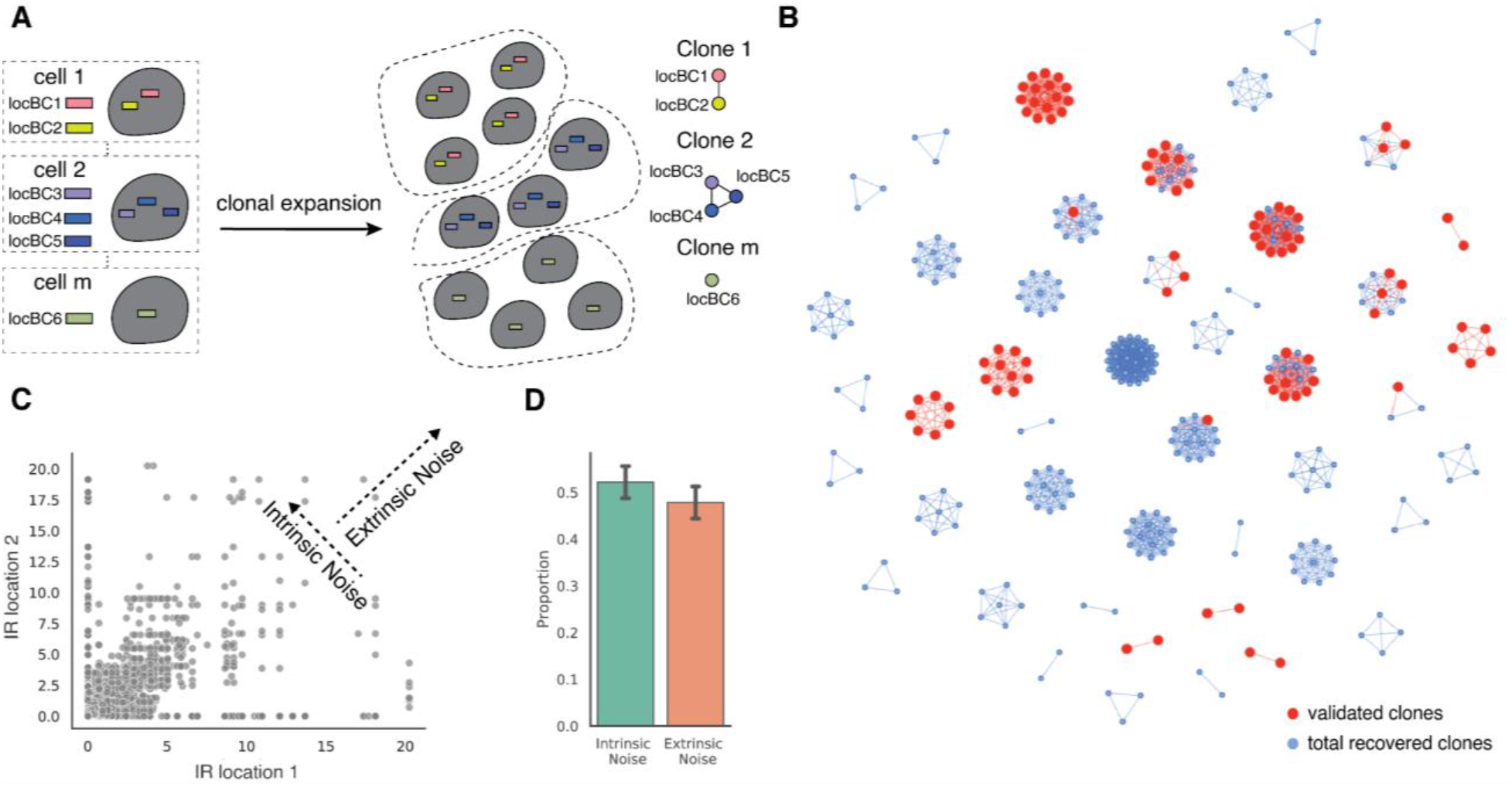
SARGENT quantifies extrinsic portion of expression noise. **(A**) Schematic for identifying different initial clones. **(B)** A network representation of the different clones identified, red nodes indicate IR locations that were independently validated by sequencing individual clones. **(C)** Pairwise expression for any two IR locations observed in the same cell. The trend along the diagonal suggests the existence of extrinsic noise, and the anti-correlation indicates the amount of intrinsic noise. **(D)** Quantification of intrinsic and extrinsic proportion of noise. Error bars from two technical replicates.

We next asked if extrinsic factors also contribute to the observed gene expression noise. For each cell in a clone, we calculated the standard deviation relative to the mean of all IRs in that cell, which we define as the fluctuation index. Lower fluctuation indices indicate that the IRs in a clone fluctuate in sync (high extrinsic noise), while higher fluctuation indexes indicate that each IR varies independently (high intrinsic noise). To simulate intrinsic noise, we first shuffled the cell labels of all the IRs within a clone and computed a distribution of fluctuation indexes for the shuffled population. If all the measured noise was intrinsic, then the measured distribution would perfectly overlap the shuffled distribution. If all the measured noise was extrinsic, then all the cells would have fluctuation indexes of 0 **(Supplementary Figure 4B)**. We found that all clones show a distribution of fluctuation indexes that is lower than that of the shuffled distribution and above zero **(Supplementary Figure 4C)**. This suggests that some portion of the expression noise can be explained by extrinsic factors that impact all IRs within a cell in different genomic environments.

To quantify the contribution of intrinsic and extrinsic noise in each clone we employed an established statistical framework^36^. Using the pairwise IR single cell expressions for all clones that contain more than one IR as input, we found that intrinsic noise comprises approximately 54% of the total noise **(Figures 4C, 4D)**. This analysis suggests that both the intrinsic chromatin context and extrinsic cellular context explains about half of the total noise in each clone. These results show that SARGENT can quantify both intrinsic and extrinsic contributions to expression noise.

### Cell substates are a source of expression noise

What cellular mechanisms control expression noise? We hypothesized that differences between cellular substates within isogenic populations are an important source of noise. Isogenic K562 cells transition between “stem-like” and “more differentiated” substates^37,38^. The stem-like substate is marked by high CD24 expression and proliferates at a higher rate, which we hypothesized would contribute to extrinsic noise. This hypothesis predicts that the same IRs will have higher MIN in stem-like cells compared to more differentiated cells. To test this prediction we sequenced the single-cell transcriptomes associated with 356 of the 939 genomic locations in parallel with the IRs. Using the transcriptomes we identified clusters of cells with high CD24 expression and confirmed that these clusters had the signatures of high-proliferating cells (**Supplementary Figures 5A, 5B**). We then calculated the expression mean and MIN for each IR location separately in the two substates. IR locations in the stem-like substate have higher mean and lower MIN **(Figures 5A, 5B)** suggesting that the global differences between the two substates are a source of MIN.

**Figure 5:**
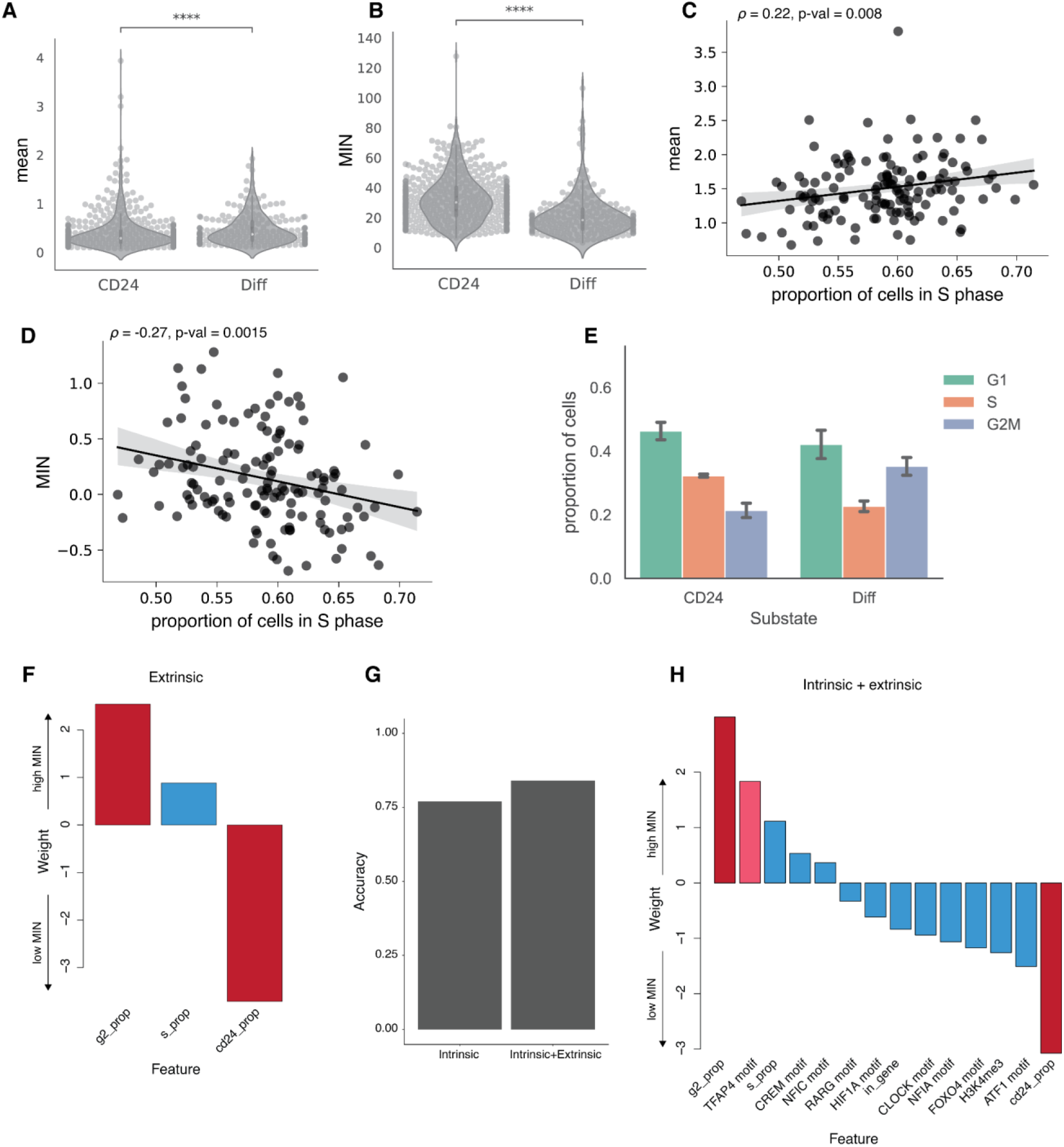
Cellular information improves classification of low vs high MIN IR locations. **(A**,**B)** Violin plots of expression mean and MIN at two substates (student t-test, ****: p < 0.0001), each dot is an IR location. (**C**,**D**) scatterplots of proportion of cells in the “stem-like” substate against mean and MIN, each dot is the average mean expression or MIN from a clone. Line: linear fit with 95% CI. Spearman correlation between mean and proportion of cells in the “stem-like” substate: 0.22, p-value = 0.08. **(E)** Barplot of the fraction of cells in different cell cycle phases for cells in the “stem-like” substate and the “differentiated” substate (Binomial test: S phase p<2.2e-16, G1 phase p <5.9e-5, G2M phase p <2.2e-16). The error bars are derived from the two replicates. **(F)** Weights of logistic regression model using extrinsic (cellular) features alone. **(G)** Addition of extrinsic features helps to improve the accuracy of the model. **(H)** Weights of logistic regression model using both intrinsic and extrinsic features. The most significant features are still the proportion of cells in the G2 phase and CD24^+^ phase. Red bars: p-value < 0.05; Pink bars: 0.05 < p-value < 0.1 from the logistic regression model.

Given the differences in mean and MIN between the substates, the MIN of the IR locations in a given clone should be partly explained by the proportion of its cells in each substate. Consistent with this prediction, we found that clones with a higher proportion of cells in the stem-like substate have slightly higher average mean expression (Spearman’s *ρ* = 0.22, p-value = 0.008), and lower average MIN (Spearman’s *ρ* = -0.27, p-value = 0.0015) across all IRs in the clone **(Figures 5C, 5D)**. We hypothesized that this was due to the slightly higher proliferation rates of cells in the stem-like phase. As expected, there are more cells in the S phase in the stem-like substate compared to the more differentiated state **(Figure 5E)**. We then examined the differences of mean and MIN in different cell cycle phases and found that expression mean is higher and MIN is lower in the S phase compared to other phases **(Supplementary Figure 5C, 5D)**. These results suggest that differences in proliferation rates is an important source of extrinsic noise, and that SARGENT is a powerful tool to dissect the extrinsic sources of expression noise.

### Cellular information improves classification of low vs high MIN IR locations

Since extrinsic factors play an important role in determining expression noise, we trained a logistic regression model to predict MIN using three extrinsic features (proportion of cells in S, proportion of cells in G2, and proportion of CD24^+^ cells). Using only the global features, the model achieved 75% accuracy (albeit without a holdout set to test on due to the small numbers of locations with associated extrinsic features). This result implies that these cellular features explain a significant portion of the variance in MIN between high and low IR locations. The proportion of cells in G2 and the proportion of cells in the CD24^+^ state were significant predictors in this model **(Supplementary Table 2)**. Being in G2 increases the probability of a high MIN IR location whereas having a higher proportion of CD24 cells reduced the probability of being a high MIN IR location **(Figure 5F)**. When we combined the significant intrinsic features from the previous model with these extrinsic features, the model accuracy increased to 84% **(Figure 5G)**. In the combined model, the extrinsic features have higher weights than the intrinsic genomic environment features **(Figure 5H)**, suggesting that the cell-state information may play a larger role in regulating MIN compared to genomic environments.

We observed a similar role for extrinsic features in classifying IR locations with high mean levels from IR locations with low mean levels. The model accuracy for just the extrinsic feature model is 80% and increases to 89% for the combined model with both intrinsic and extrinsic features **(Supplementary Figure 5E)**. In the combined model, the proportion of cells in the CD24 cell-state is the most highly weighted feature **(Supplementary Figure 5F, Supplementary Table 3**). In contrast to the MIN model, the proportion of cells in the CD24 state increases the probability of being a high-mean IR location **(Figure 5H, Supplementary Figure 5F**), which is consistent with our observations in **Figures 5B and 5D**. Thus, while cellular information plays an important role in gene expression regulation, these features have orthogonal impacts on expression mean and single-cell variability.

### Effects of transgenes integration on endogenous genes

Finally, SARGENT can be used for purposes beyond studying gene expression noise. One such application is screening for ‘safe harbor’ loci in the genome. To achieve safe and effective gene therapy, we need to identify genomic locations that have stable expression of the transgene of interest (high mean expression and low noise) and have minimal effects on endogenous gene expression. Historically, transgenes are often integrated into several known ‘safe harbor’ loci^39^. Those loci are mainly located in the introns of stably expressed genes to prevent silencing. Because SARGENT can be used to measure gene expression mean, noise and endogenous gene expression simultaneously, we can leverage SARGENT to screen for potential safe harbors in a high-throughput manner.

We examined how our reporter gene integrations altered the expression of the gene into which it integrated. We focused on the 65 IR locations that are integrated into gene bodies **(Supplementary Table 5)**. These integrations were distributed across different clones **(Supplementary Figure 6A)** and should not be confounded by clonal effects. We calculated pseudo-bulk expression for each gene from clones that contain the integration and compared that to the expression from other clones that do not have the IR integration **(Figure 6A)**. We found that in most cases (61/65), transgene integration does not alter the endogenous gene expression **(Figure 6B)**. We also randomly shuffled the gene labels to compute the background differential expression, and found that there were no significantly differentially expressed genes once the labels were shuffled **(Supplementary Figure 6B)**. Among the locations with significantly differentially expressed genes, 3 out of 4 IR integrations increases gene expression **(Figure 6C)**, consistent with previous studies showing that the integration of a transgene often increases endogenous gene expression^41^. Taken together, our results suggest that most endogenous genes are not impacted by the integration of exogenous genes. This result illustrates that SARGENT could be a powerful tool to screen for “safe harbor” loci for transgene integration.

**Figure 6:**
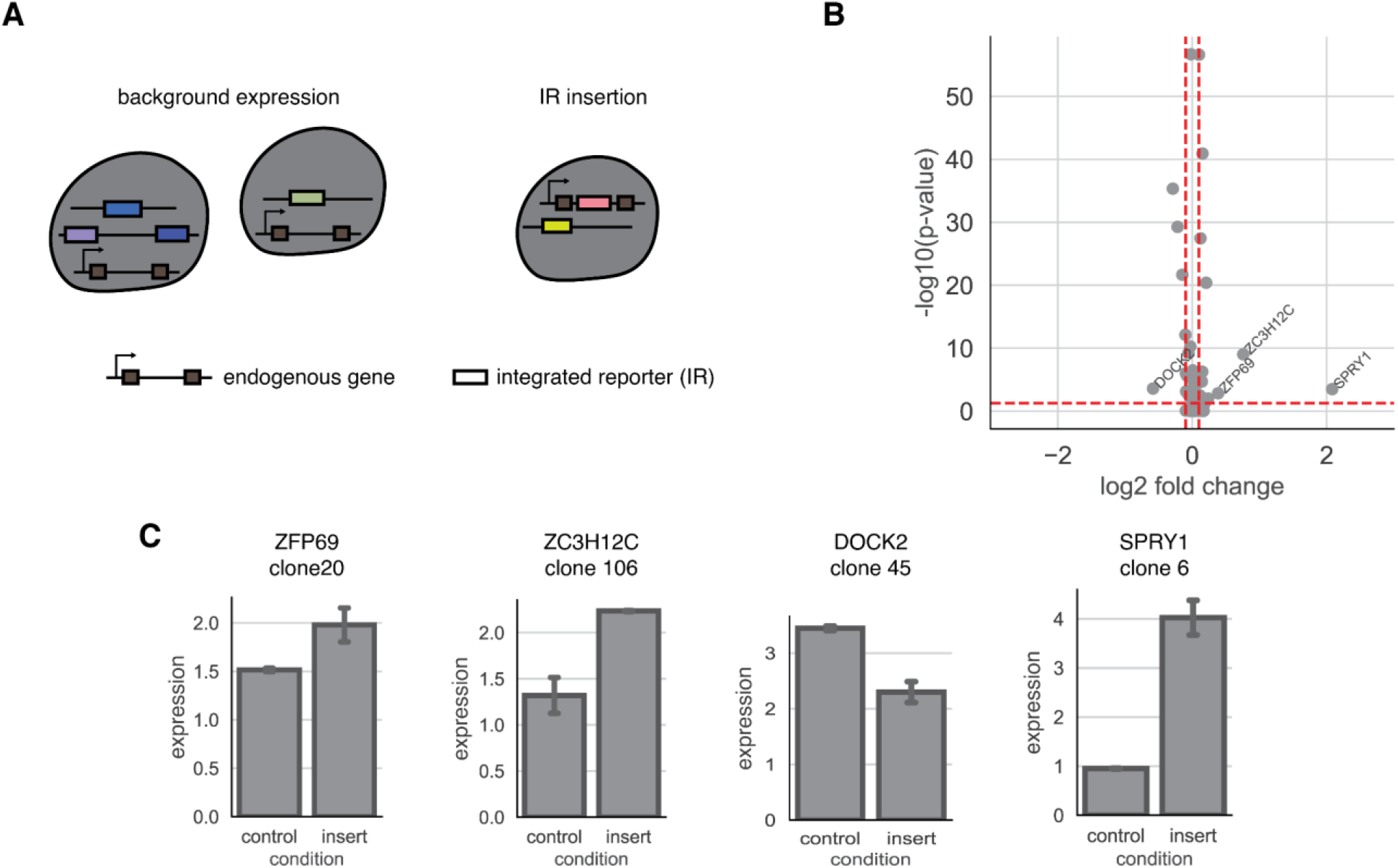
SARGENT measures the insertion effect of a transgene. (**A**) Schematic for expression change detection in the transcriptome data. (**B**) Volcano plot of log2 fold change and -log10(p-value) from a Fisher’s Exact Test. Red dotted line: cut off for fold change (0.5), cut off for p-value: 0.05. (**C**) Barplots of difference of expression between genes without IRs (control) and genes with IRs (insert). The clone where the IR is integrated is indicated. Error bars are derived from two technical replicates. < p-value < 0.1 from the logistic regression model.

## Discussion

Since the early single-cell studies showing the variability of gene expression in isogenic populations^25^, many individual chromatin and sequence features have been suggested to modulate expression noise^1,5,42,43^. However, there has yet to be a systematic study of the impact of different genomic features on large numbers of identical genes.

We developed SARGENT, a high throughput method to measure the expression mean and noise at different genomic locations in parallel. One key advantage of SARGENT is that the reporter gene used in all locations is identical, which allows us to isolate the effects of the genomic environments without being confounded by the effects of different promoters. We identified different chromatin marks that are associated with high or low MIN, and used a logistic regression model to identify features of the genomic environments that might control MIN. Our observations indicate that the features that control expression noise are independent of the features controlling expression mean. Several recent studies have developed tools for the orthogonal control of mean and gene expression noise^42,44,45^. To this end, our results suggest potential mechanisms that can be targeted for independent modulation of expression mean and single-cell variability.

We also quantified the extrinsic portion of expression noise and identified that the oscillation between a “stem-like” substate and a “differentiated” substate in K562 cells is an important source of extrinsic noise. Our data suggests that extrinsic noise might be more important in regulating MIN than genomic environments. This indicates that the regulation of noise of individual genes might be at the level of the promoter, rather than through its chromatin or genomic environment.

We envision that SARGENT will be a useful tool for other synthetic biology applications. While advances in genome engineering technologies now allow researchers to integrate transgenes at most desired genomic locations, the selection of appropriate sites for transgene overexpression remains non-trivial, with no location in human cells validated as a safe harbor locus^41,46^. This is mainly due to the lack of methods to systematically screen for loci that have high expression, low variability and do not impact cellular function. Here we showed that SARGENT can be used to read out a transgene’s impact on global expression as well as the endogenous gene that it is integrated into. Surprisingly, most endogenous genes are not impacted by the insertion of reporter genes. With SARGENT, we can quickly screen genomic locations to find the best locations for human transgene integration which will prove useful gene therapy applications.

More broadly, we envision that SARGENT will be a useful technology for many different applications including mechanistic studies of gene expression noise and synthetic biology applications. The 10x Genomics platform used in this study is limited by throughput, but improvements to scRNA-seq technologies will increase the scope of SARGENT. For example, coupling sci-RNA-seq^47^ or SPLiT-seq^48^ to SARGENT would allow for many more locations to be assayed in parallel. A larger goal will be to construct a detailed map of the MIN landscape across the genome, much like the maps of mean expression levels generated by ENCODE.

## Methods

### SARGENT library cloning

All primers and oligonucleotides used in this study are listed in **Supplementary Table 6**. To clone the reporter gene for SARGENT, we first cloned a CMV-BFP reporter gene containing the 10x capture sequence 1 (CS1) into a piggyBac vector containing two parts of a split-GFP reporter gene^49^. When the reporter gene construct is integrated into the genome, the split-GFP combines to produce functional GFP, allowing us to sort for cells that have successful reporter gene integrations. We next added a library of random barcodes to the plasmid by digesting the plasmid with XbaI followed by HiFi assembly (New England Biolabs) with a single-stranded oligo containing 16 random N’s (Location barcodes; locBC) and homology arms to the plasmid (CAS P57).

### Generation of cell lines for SARGENT

K562 cells were maintained in Iscove’s Modified Dulbecco′s Medium (IMDM) + 10% FBS + 1% non-essential amino acids + 1% penicillin/streptomycin. We selected two K562 cell lines previously used in our lab that each contain a ‘landing pad’ at a unique location with a pair of asymmetric Lox sites for recombination (loc1 - chr8:144,796,786, loc2 - chr11: 16,237,204; hg38 coordinates). Using these ‘landing pad’ cell lines allows us to perform smFISH on the landing pad to directly compare SARGENT and smFISH results. For each cell line, we replaced the original landing pad cassette with the same reporter gene in the SARGENT library so that we can capture the reporters from the landing pad and reporters from other genomic locations in SARGENT using the same primers. Pool 1 was derived from the loc2 cell line, while Pools 2, 3 and 4 were derived from the loc1 cell line.

The SARGENT library and piggyBac transposase were co-transfected into K562 (LP cell lines) cells at a 3:1 ratio using the Neon Transfection System (Life Technologies). For each experiment, we transfected 2.4 million cells with 9μg of SARGENT library and 3μg of transposase. The cells were sorted after 24 hours for GFP-positive cells to enrich for cells that have integrated SARGENT reporters. We reasoned that ∼100 single cells for each Integrated Reporter (IR) location would be required to obtain a good estimate of mean and variance. Each SARGENT experiment contains many single cell clone expansions: all the cells from the same clone share the same genomic integrations. Since we targeted approximately 20,000 cells per 10x run, the upper limit of the numbers of clones we can test in one experiment is 200. Because 10x also has a high dropout rate, we targeted 100 clones per experiment in order to ensure that we obtained high quality data. Each clone has an average of 5 integrations, which theoretically allows us to assay 500 IR locations in one experiment. Since the clones did not all grow at the same rate, practically we obtained fewer than 500 IRs per experiment.

For Pools 1 and 2, cells were sorted into pools of 100 cells each and allowed to grow until there were sufficient cells for RNA/DNA extraction and SARGENT experiments. Pool 3 contained the same cells as Pool 2, except that single cells were allowed to grow individually and pooled by hand just before the SARGENT experiments. This allowed for a more even representation of each individual clone (which contains unique integrations) in the final pool. For Pool 4, transfected cells were first sorted into 96-well plates with 2 cells/well and allowed to grow individually and 100 wells were manually pooled for SARGENT experiments. We used cells from Pool 4 to compute technical reproducibility.

### SARGENT integration mapping

We harvested DNA from SARGENT pools using the TRIzol reagent (Life Technologies). To map the locations of SARGENT integrations, we digested gDNA for each pool with a combination of AvrII, NheI, SpeI and XbaI for 16 hours. The digestions were purified and self-ligated at 16°C for another 16 hours. After purifying the ligations, we performed inverse PCR to amplify the barcodes with the associated genomic DNA region (CAS P59 and P64). For each pool, we performed 2 technical replicates with 8 PCRs per replicate and pooled the PCRs of each replicate for purification. We then used 8ng of each replicate for further amplification with 2 rounds of PCR to add Illumina sequencing adapters (CAS P55 and P65). The sequencing library was sequenced on the Illumina NextSeq platform.

The barcodes of each read were matched with the sequence of its integration site. The integration site sequences were then aligned to hg38 using BWA^50^ with default parameters. Only barcodes that mapped to a unique location were kept for downstream analyses. All barcodes and IR locations can be found in **Supplementary Table 1**.

### ClampFISH

Single-molecule FISH was performed on the two ‘landing pad’ locations that were in the original cell lines used for SARGENT (see Generation of cell lines for SARGENT above). ClampFISH probes for the reporter genes were designed using the Raj Lab Probe Design Tool (rajlab.seas.upenn.edu, **Supplementary Table 7**). Each probe was broken into three arms to be synthesized by IDT. The 5’ of the left arm is labeled by a hexynyl group, and the 3’ of the right arm is labeled by NHS-azide. The right arm fragment was purified by HPLC. All three components were resuspended in nuclease-free H2O to a concentration of 400 uM. The three arms were ligated by T7 ligase (NEB, Cat# M0318L), at 25 C overnight. then purified using the Monarch PCR and DNA cleanup Kit (NEB, Cat# T1030S) and eluted with 40 ul of nuclease-free water. After the ligation, each probe is stored at -20 C. ClampFISH was performed according to the suspension cell line protocol of clampFISH^51^. 0.7 million cells were collected and fixed in 2 mL of fixing buffer containing 4% formaldehyde for 10 min, then permeabilized in 70% EtOH at 4 C for 24 hours. The primary ClampFISH probes were then hybridized for 4 hours at 37 C in the hybridization buffer (10% Dextran Sulfate, 10% Formamide, 2X SSC, 0.25% Triton X). After hybridization, cells were spun down gently at 1000 rcf for 2 min. Cells were washed twice with the washing buffer (20% formamide, 2X SSC, 0.25% Triton X) for 30 min at 37 C. The secondary probes were then hybridized to cells at 37 C for 2 hours and the cells were then washed twice with washing buffer for 30 min at 37 C. The primary and secondary probes are “clamped” in place through a click reaction (CuSO4 75 uM, BTTAA 150 uM, Sodium Ascorbate 2.5 mM in 2X SSC) for 20 min at 37 C. The cells were then washed twice in the washing buffer at 37C for 30 min each wash. Then, the cells were hybridized with the hybridization buffer with tertiary probes for 2 hours at 37C. We complete 6 cycles of hybridization for all our experiments. After the final washes, cells were incubated at 37 C with 100mM DAPI for 20 min, washed twice with PBS, resuspended in the anti-fade buffer, and spun onto a #1.5 coverslip (part number) using a cytospin cytocentrifuge (Thermo Scientific), mounted onto a glass slide, sealed with a sealant, and stored at 4C.

### SARGENT library using the 10x Genomics platform

#### Cell preparation

We used the Chromium Single Cell 3’ Kit (v3.1) from 10x Genomics for SARGENT. We followed the manufacturer’s instructions for preparing single-cell suspensions. We used a cell counter to measure the number of cells and viability, and used cell preparations with greater than 95% cell viability.

#### Cell barcoding and reverse transcription

We followed the manufacturer’s instructions with the following modifications in Pools 1-3: no 10x template switching oligo (PN3000228) was added to the Master Mix (Step 1.1). To correct for the missing volume, 2.4 μl of H_2_O was added to the master mix per reaction. For pool 4, the template switching oligo was included as written. For the cDNA amplification (Step 2.2), no 10x provided reagents were used. Instead, a custom primer (CAS P20) was used with 14 cycles of amplification with the provided 10x protocol (Step 2.2 d). For the pool where we also sequenced transcriptomes (Pool 4), we followed the 10x protocol as written for cDNA amplification.

#### Barcode PCR and library preparation

We performed nested PCRs to amplify barcodes from 10x cDNA. For pools 1-2, PCR library construction was split into two pools for amplification of transcripts captured by capture sequence 1 and poly(A) respectively. Both PCR reactions were done with 2μl purified cDNA, 2.5μl 10 μM reporter-specific forward primer (CAS P45), 2.5μl 10 uM poly(A) (CAS P20) or capture sequence adapter-specific primers (CAS P32), and 25μl Q5 High Fidelity 2X Master Mix (M0492, New England Biolabs) in 50μl total volume with 10 cycles amplification. The PCRs were then purified with Monarch PCR & DNA Cleanup Kit (New England Biolabs, T1030) and Illumina adapters were added in another 2 rounds of PCR, with a PCR purification step with the Monarch kit between PCRs. For poly(A) amplicons, we used CAS P42 and CAS PP2, followed by CAS P48 and CAS PP4. For capture sequence amplicons, we used CAS P41 and CAS CS2, followed by CAS P48 and CAS CS4. The reactions were then pooled and purified with SPRIselect Beads (Beckman Coulter) at 0.65x volume. For pool 4, we performed the PCRs for the poly(A) fraction using 2μl purified cDNA as described above, but not the capture sequence transcripts.

### SARGENT data processing

#### Read parsing

We first identified the reads that match the constant sequence in our reporter gene. We used two versions of constant sequence to match against, depending on if the read was captured using the poly(A) sequence on the mRNA or the capture sequence specific to the 10x beads. We used a fuzzy match algorithm to capture reads that have a mismatch at these positions due to sequencing error. From each read, we parsed out the cell barcode, 10x UMI and locBC. We then collapsed reads with identical cell barcodes, UMI and locBCs into one “trio” and kept track of the number of reads supporting each trio. For downstream analysis, we filtered out trios with low numbers of supporting reads since these are likely to be enriched for PCR artifacts. We next processed the trios to error correct the cell barcodes and locBCs before estimating the mean and variance.

#### Barcode error correction

To correct for PCR artifact and sequencing errors, a custom script was used to error-correct for 10x cell barcodes. Briefly, we first acquired the empirical distribution of the Hamming distances among observed 10x cell barcodes. We found that more than 99% of 10x cell barcodes are more than 6 hamming distances away from each other, making error correction a feasible approach to denoise the data. We first identify cell barcodes that match perfectly to the 10x cell barcode whitelist, then we order them based on their abundance. The cell barcodes that are not in the whitelist are then compared to the ordered whitelisted cell barcodes, if the Hamming distance between the non-whitelisted cell barcodes is within 2 Hamming distances of a whitelisted cell barcodes, we correct the unwhitelisted cell barcode. With cell barcode correction, we recovered ∼12% of reads that would have been discarded.

Due to the random synthesis of the locBC, a slightly different approach was taken for error correction for the locBCs. Briefly, all the locBCs are ranked based on abundance. Starting from the most abundant barcode, we look for locBCs that are within 4 Hamming distance to that barcode and correct them. We then remove that barcode and any corrected barcodes, and repeat this process until we have iterated through all locBCs.

#### Calculating mean and variance of each IR

We filtered out cells that had less than 5 IR integrations (locBCs) and less than 10 UMIs. We also filtered out locBCs that were seen in less than 5 cells and UMIs that had less than 2 supporting reads. We then computed the number of UMIs per locBC in each cell to calculate the expression level of each locBC. For each locBC, mean expression was calculated as the average UMI count across all cells that expressed that locBC. Expression variance was calculated as the variance in UMI counts across all cells that expressed that locBC.

#### Mean-independent noise (MIN) metric

In order to remove the effect of the mean on the variance we first fit a linear model: log2(variance of IR location) ∼ log2(mean of IR location) for each experimental pool and used the residuals of the model as the mean-independent noise metric. For each IR location, the MIN is the residual variance after removing the effect of the mean.

### Analyses of genomic environment effects on mean-independent noise

#### Chromatin environment association with mean/MIN

We downloaded the Core 15-state chromHMM annotations for K562 cells from the Roadmap Epigenomics Project^22^. We then collapsed similar annotations and overlapped the IR locations with the corresponding annotation using the GenomicRanges R package^52^.

We split the IRs into locations with high vs low mean or high vs low MIN respectively. We then downloaded histone ChIP-seq datasets from ENCODE^35^ **(Supplementary Table 8)** and plotted the signals 10kb surrounding each class of IRs using the ComplexHeatmap package in R^53^.

To look for enriched TF motifs we first downloaded all human motifs from the HOCOMOCO v11 database. We then filtered the motifs for TFs that are expressed (FPKM ≥1) in the K562 cell line using whole-cell long poly(A) RNA-seq data generated by ENCODE (downloaded from the EMBL-EBI Expression Atlas, **Supplementary Table 8**). We then used the STREME package^54^ (MEME suite 5.4.1) with sequences of 1kb surrounding each IR to identify enriched *de novo* motifs in high or low MIN regions, using the other class as the control set of sequences. We then took the top 2 motifs for each bin and matched it against a list of TFs expressed in K562s using TOMTOM^55^ (MEME suite 5.4.1). We reported the top 6 TOMTOM matches.

#### K562 Hi-C

We performed Hi-C on wild-type K562 cells with the Arima Hi-C kit (A510008) according to the manufacturer’s protocols (3 replicates, 870 million reads total). The reads were then processed with the Juicer pipeline^56^ to generate HiC contact files for each replicate. We then used the peakHiC tool^57^ to call loops from each IR with the following parameters: window size = 80, alphaFDR = 0.5, minimum distance = 10kb, qWr = 1. Using these parameters each IR was looped to a median of 3 regions (range 0-7).

#### Logistic Regression model for Intrinsic and Extrinsic features associated with MIN

We used histone ChIP-seq and ATAC-seq datasets from ENCODE^35^ **(Supplementary Table 8)** and overlapped their signals with each IR using used bedtools v2.27.1^58^. For all features we considered the 20kb upstream and downstream of each IR respectively. For each histone modification, we computed the mean ChIP signal around the IRs. For ATAC-seq, we calculated the total number of peaks with the bedtools map count option. To look for TF motifs we counted the numbers of each motif for TFs expressed in K562s (see above) in each surrounding IR sequence using FIMO^59^ (MEME suite 5.0.4). Because this resulted in a long list of TFs we further filtered the TFs to include only those with a significant correlation with MIN levels in the regression model. To determine the numbers of enhancers interacting with each IR we annotated the loops called from peakHiC above with chromHMM enhancer annotations using the GenomicInteractions R package^60^ and counted the number of enhancers.

For the extrinsic features, we calculated the proportion of cells in the “stem-like” substate and “differentiated” substate and different cell cycle phases based on the barcodes that appeared in those substates. We removed IR locations that have less than 30 cells in any of the substates.

We used the glm function in R (version 3.6.3) to fit logistic regression models. We separated the IR locations into top 20% MIN and bottom 20% MIN and used logistic regression to classify locations. We first fit a model with just local sequence features (chromatin modifications, number of TF motifs, number of loops, whether the IR location is in a gene, GC content and the number of ATAC-seq peaks). We used data from one experiment for training the model and used data from another experiment as a holdout set of data to estimate the performance of the classifier. We next fit a model with cellular information for each IR location: proportion of cells with data for the IR location in S phase of the cell cycle, in G2 phase and the proportion of cells that are in the “stem-like” substate of K562 cells^37^. Lastly, we fit a model that incorporated the extrinsic features and the significant predictors from the intrinsic features model.

### Transcriptome analyses associated with SARGENT

#### Processing the single-cell transcriptome data

The single-cell RNAseq data was processed with CellRanger 6.0.1 and Scanpy 1.9.1. Briefly, the raw reads were processed with the standard single-cell expression cell line pipeline line. The resulting expression matrix was then imported into Scanpy for further visualization and clustering.

#### Identifying single cell clones

We identified the individual clones for Pool 4 which contained cells that grew out of 100 two-cell clones. Since most of the clones will have unique integrations into unique genomic locations, the cells that grew out from the same clone will have identical unique sets of locBCs. Due to the dropout rates associated with scRNAseq methods, not all barcodes will be present in all cells, nor will the cell barcodes be uniquely linked to correct sets of locBCs. To identify the barcodes belonging to the same clone, we first recorded locBCs that are linked by a given cell barcode. We then filtered the locBC list associated with a given cellBC based on the number of UMIs associated with these locBC. At this step, we used a knee point detection algorithm^61^ that automatically detects the inflection point of the ordered UMI counts histogram. After filtering for locBCs that appear in more than 5 cells, we constructed a clonal graph by linking locBCs that co-occur in the same cells.

#### Validation of individual clones

We extracted gDNA from 16 clones that were grown out from Pool 4. We then amplified the barcodes from each clone using Q5 High Fidelity 2X Master Mix (M0492, New England Biolabs) with primers specific to our reporter gene (CAS P58-59). For each clone, we performed 4 PCRs and pooled the PCRs for purification. 4ng from each clone was then further amplified with 2 rounds of PCR to add Illumina sequencing adapters (CAS P60-63). The barcodes were sequenced on the Illumina NextSeq platform.

#### Estimating intrinsic vs extrinsic noise

To understand how cellular environments affect IR expression, we computed the mean and standard deviation from all IR locations in the same cell. Since standard deviation is expected to increase with mean, we calculated the standard deviation/mean for each cell, which we termed the fluctuation index **(Supplementary Table 9)**. To establish the null distributions, we randomly shuffled the cell labels for each clone and computed fluctuation indices for the shuffled cells.

Intrinsic and extrinsic noise were estimated using the statistical framework developed for the dual-reporter experiment^36^. In our experiment, single-cell expression differences among IR locations are treated as the intrinsic portion of the noise. We first extracted the pairwise expression level for IR locations in every single cell. We then applied the statistical framework developed by Fu and Pachter^36^. The derivation is abbreviated and can be found in the original publication. Briefly, let C denote the expression for the first locBC in the cell, Y denote the expression for the second locBC in the cell and n denote the number of cells.

Let ŋ_ext_ denote the extrinsic noise, and it can be calculated as:

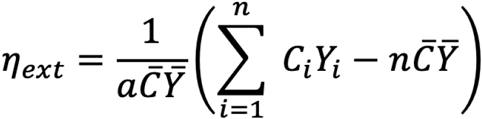

where

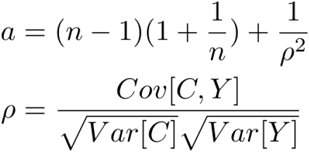

Similarly, let ŋ_int_ denote the intrinsic noise, and it can be calculated as:

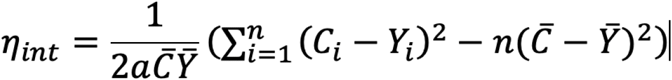

where

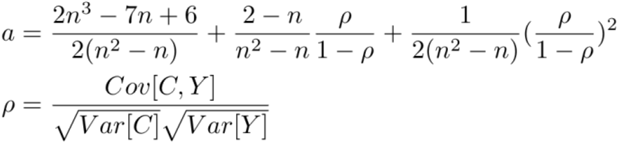

#### Cell substate impact on expression mean and noise

To compute cell substate specific expression mean and noise at different genomic locations, individual cells were assigned a cell cycle phase of G1, S, or G2/M using a previously reported set of cell-cycle specific marker genes with Scanpy 1.9.1^62^. For the stem-like substate analysis, we clustered cells based on their transcriptomes and assigned cells in the CD24 high cluster as CD24+ cells^37^. To ensure an accurate measurement of expression mean and noise, genomic locations with less than 15 cells in any phase were excluded from the cell cycle analysis. Based on this filtering criterion, 345 out of 939 genomic locations were used for this analysis. To determine the impact of cellular substates on gene expression noise, we calculated the proportion of cells in different cellular substates for each clone. For each clone, we also calculated the average mean and variance of all the IRs in that clone.

#### Transgene integration analysis

To examine whether the integration of a trans-gene alters endogenous gene expression, we first identified IR locations that were integrated into a gene body. Since the IR insertion only occurs in a single clone, we computed pseudobulk expression from cells in the clone using decouplerR 1.1.0^40^. We then randomly sampled the same number of cells from all the other clones and used the pseudobulk expression from these cells as wild-type expression. To determine whether the expression in the IR clone is significantly different from wild-type expression, we computed the p-value of differential expression using Fisher’s exact test.

## Acknowledgements

We thank the members of the Cohen Lab for their helpful comments and critical feedback on the manuscript. We are also grateful to Jessica Hoisington-Lopez and MariaLynn Crosby in the DNA Sequencing Innovation Lab for assistance with high-throughput sequencing; the Genome Engineering and iPSC Center for kindly allowing us to use their flow cytometer for cell sorting and the Hope Center DNA/RNA Purification Core at Washington University School of Medicine for helping with gDNA extractions.

## Author contributions

A.R, S.Z, C.K.Y.H and B.A.C conceived and designed the project. S.Z, A.R, C.K.Y.H designed and conducted all experiments and analyses. All authors wrote and edited the manuscript. C.K.Y.H, A.R, S.Z contributed equally to this project.

## Supplementary Figures

**Supplementary Figure 1:**
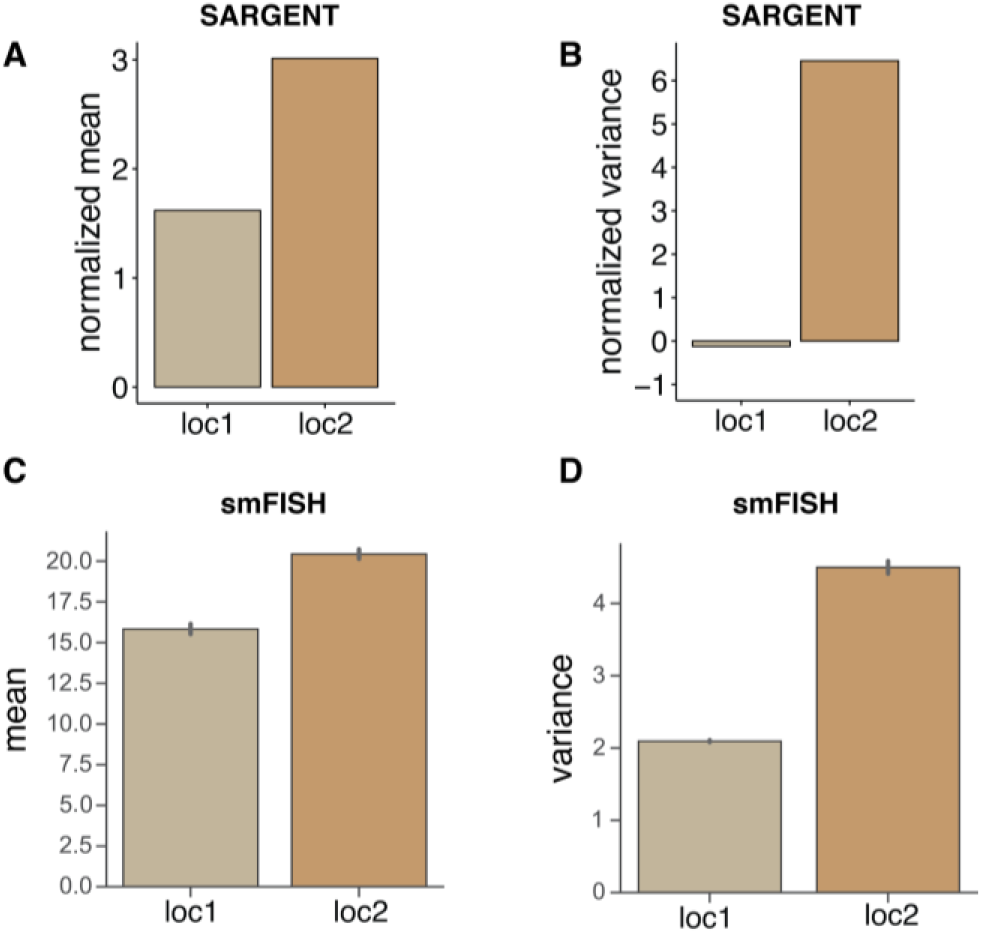
**(A, B)** Mean and noise levels of two IR locations (loc1 and loc2) measured by SARGENT. Values were normalized (z-scored) for comparison across different experiments. **(C**,**D)** Mean and noise levels of the same two IR locations measured with smFISH. Error bars represent 1 std from two biological replicates.

**Supplementary Figure 2:**
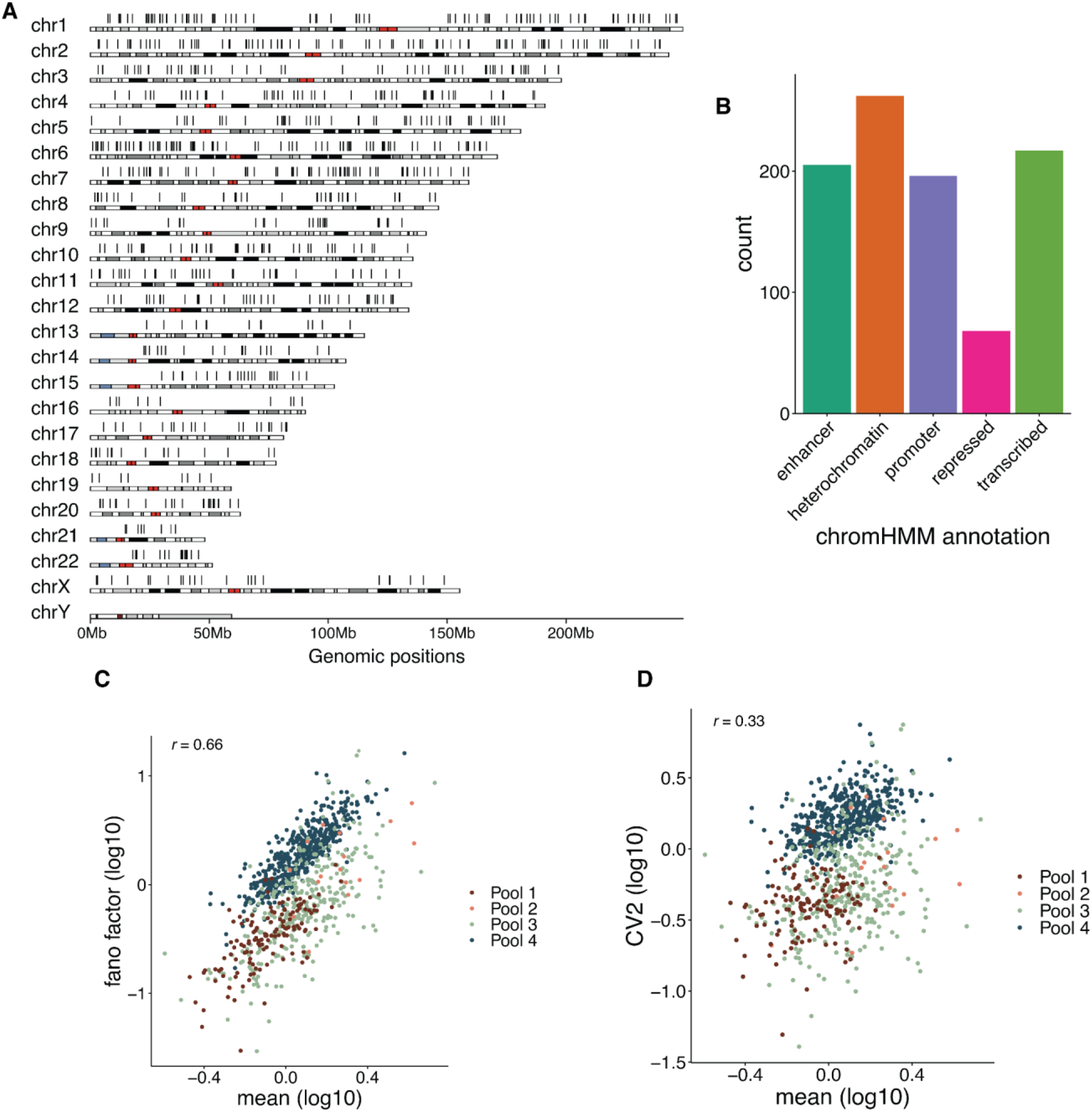
Measurements of mean-independent noise across different chromosomal environments. **(A)** IR locations are distributed all throughout the genome. Each black bar above the ideogram represents a separate integration. **(B)** IR locations are found distributed across different chromatin types. **(C, D)** Expression mean is well correlated with fano factor (C) and CV^2^ (D).

**Supplementary Figure 3:**
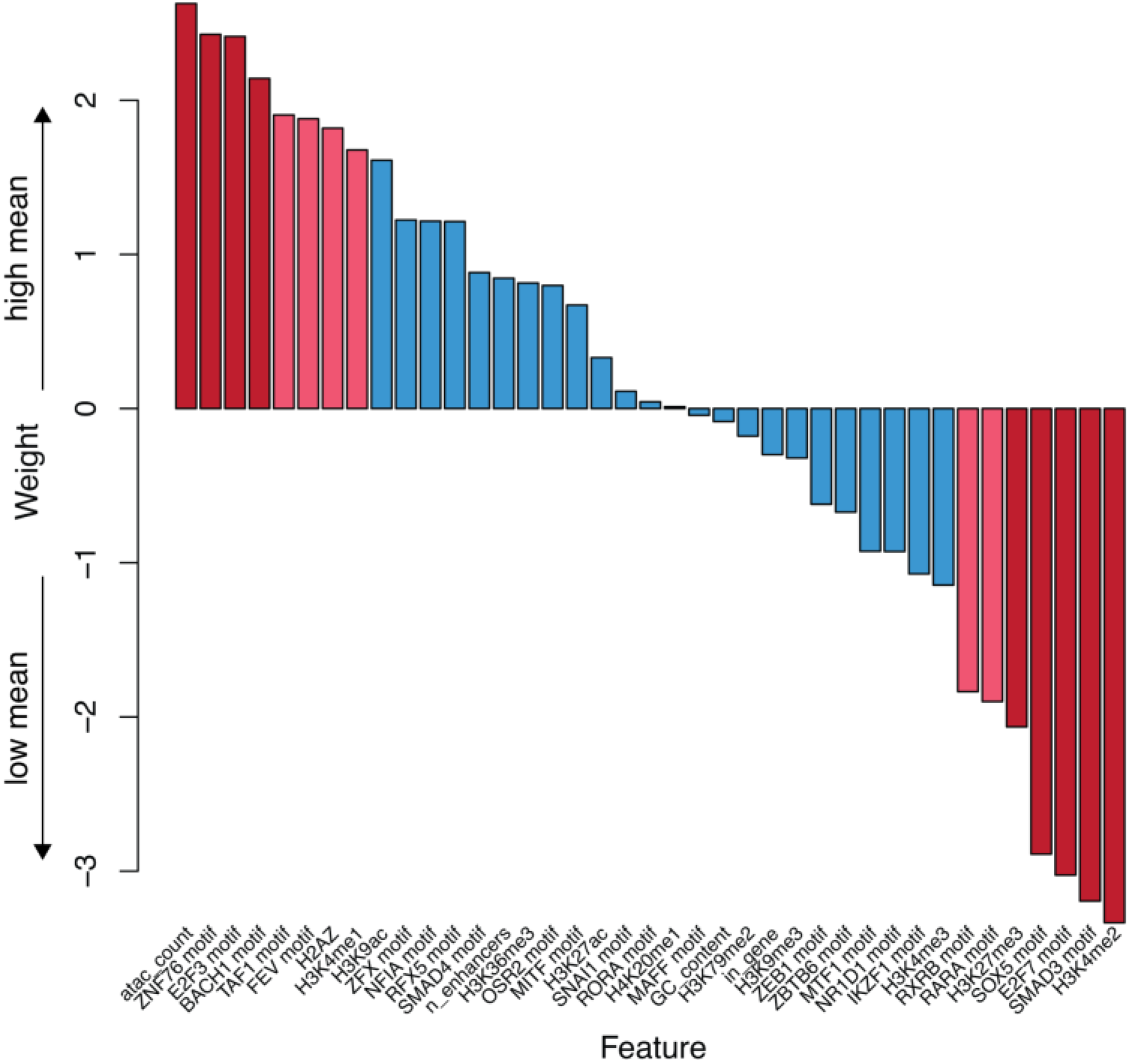
Model results for high mean vs low mean. Weights of features from the gene expression mean model using only intrinsic genomic features. Red bars: p-value < 0.05; Pink bars: 0.05 < p-value < 0.1 from the logistic regression model.

**Supplementary Figure 4:**
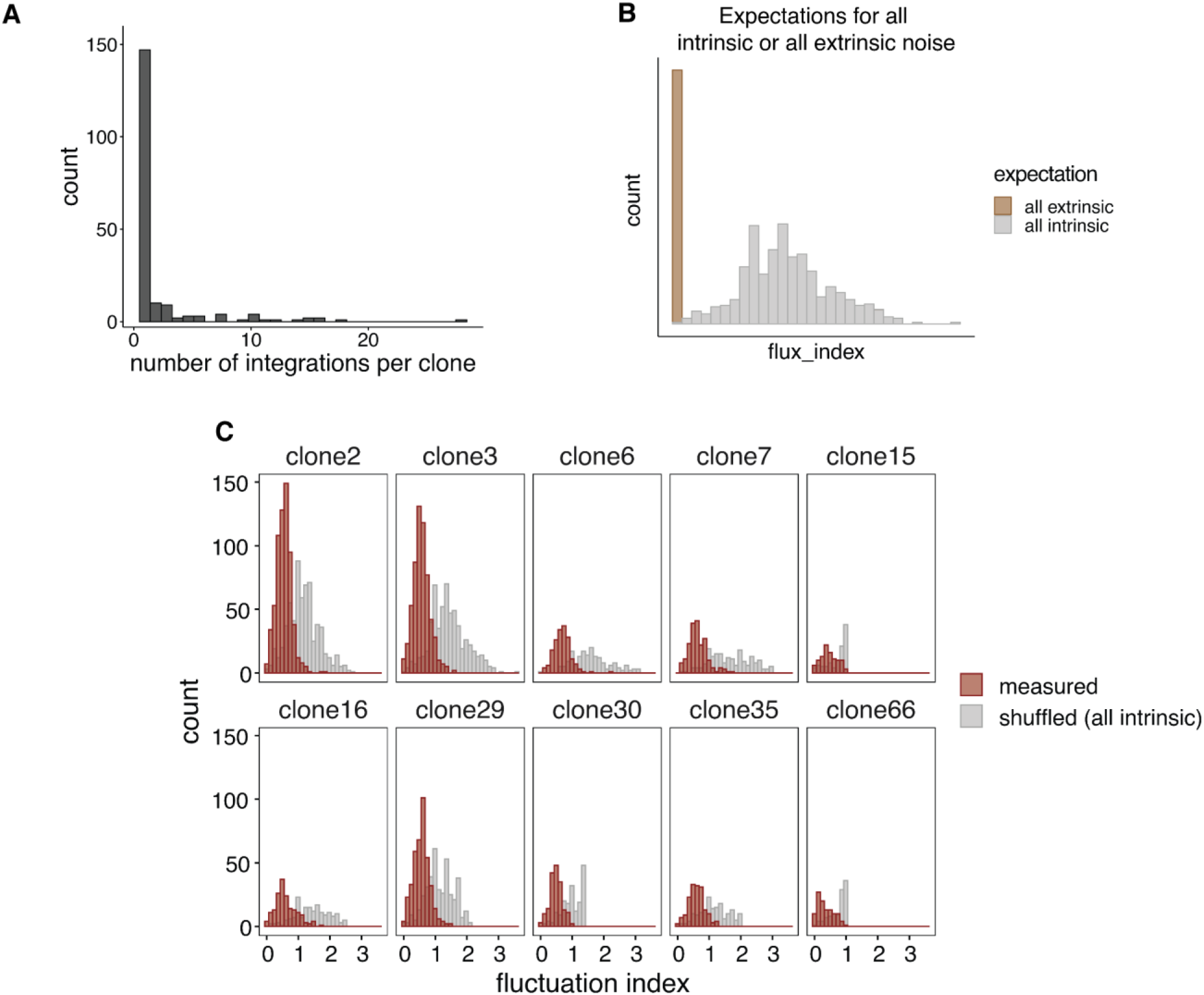
Clonal identification allows for separation of intrinsic and extrinsic noise. **(A)** Histogram of integrations per clone. **(B)** Mock histogram showing the expected distributions if noise was either all intrinsic or all extrinsic. **(c)** Histogram of measured fluctuation indices (defined as the standard deviation/mean of all IRs in a cell) for 10 random clones. The shuffled distribution represents the distribution after cell labels have been shuffled, which simulates the case when all noise is intrinsic.

**Supplementary Figure 5:**
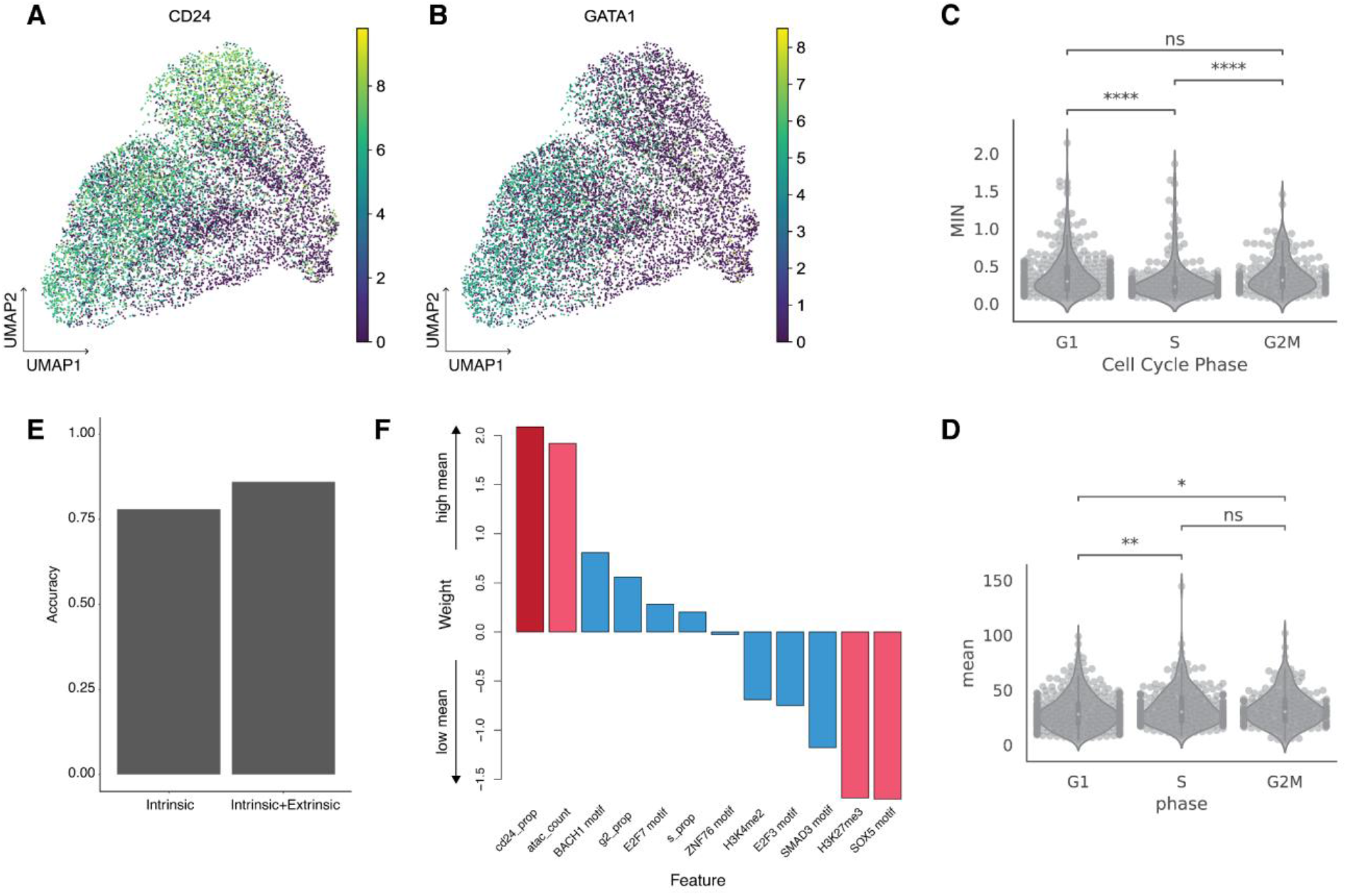
Cell cycle and CD24 states partially explain extrinsic noise. **(A)** UMAP clustering reveals CD24^+^ cell population. **(B)** CD24^+^ cells express stemness and proliferation marker gene GATA1. **(C, D)** Violin plots of MIN (C) and mean (D) levels in different phases of the cell cycle. P-values were calculated using the Mann-Whitney-Wilcoxon test. Legend: *: 0.01 < p-value ≤ 0.05, **: 0.001 < p-value ≤ 0.01, ***: 0.0001 < p-value ≤ 0.001, ****: p-value ≤ 0.0001. **(E)** Gene expression mean model improves after the addition of extrinsic features. **(F)** Weights of features from the mean model using both intrinsic genomic and extrinsic features. Red bars: p-value < 0.05; Pink bars: 0.05 < p-value < 0.1 from the logistic regression model.

**Supplementary Figure 6:**
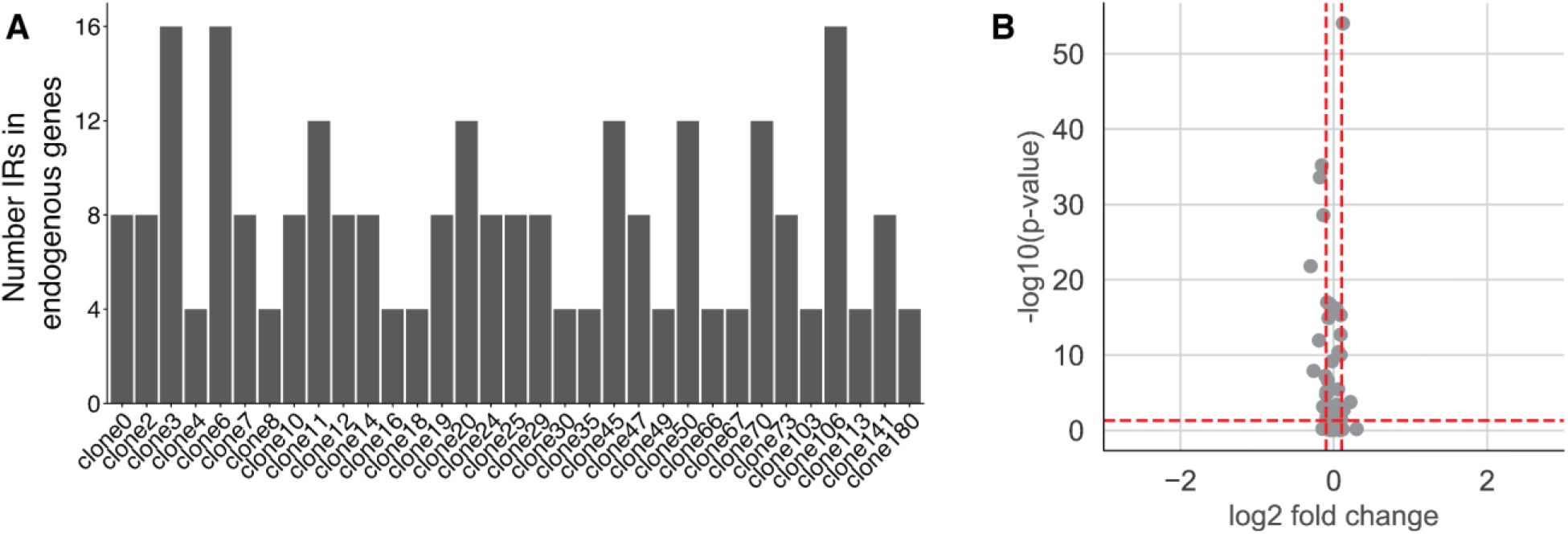
IR integrations have little impact on endogenous expression. **(A)** Bar plot number of IRs in endogenous genes per clone. **(B)** Shuffling IR-endogenous gene labels results in no differentially expressed genes.

## References

1. Raj, A. & van Oudenaarden, A. Nature, nurture, or chance: stochastic gene expression and its consequences. Cell 135, 216–226 (2008).

2. Chang, H. H., Hemberg, M., Barahona, M., Ingber, D. E. & Huang, S. Transcriptome-wide noise controls lineage choice in mammalian progenitor cells. Nature 453, 544–547 (2008).

3. Kalmar, T. et al. Regulated fluctuations in nanog expression mediate cell fate decisions in embryonic stem cells. PLoS Biol. 7, e1000149 (2009).

4. Abranches, E. et al. Stochastic NANOG fluctuations allow mouse embryonic stem cells to explore pluripotency. Development 141, 2770–2779 (2014).

5. Desai, R. V. et al. A DNA repair pathway can regulate transcriptional noise to promote cell fate transitions. Science 373, (2021).

6. Spencer, S. L., Gaudet, S., Albeck, J. G., Burke, J. M. & Sorger, P. K. Non-genetic origins of cell-to-cell variability in TRAIL-induced apoptosis. Nature 459, 428–432 (2009).

7. Topolewski, P. et al. Phenotypic variability, not noise, accounts for most of the cell-to-cell heterogeneity in IFN-γ and oncostatin M signaling responses. Sci. Signal. 15, eabd9303 (2022).

8. Weinberger, L. S., Burnett, J. C., Toettcher, J. E., Arkin, A. P. & Schaffer, D. V. Stochastic gene expression in a lentiviral positive-feedback loop: HIV-1 Tat fluctuations drive phenotypic diversity. Cell 122, 169–182 (2005).

9. Shaffer, S. M. et al. Rare cell variability and drug-induced reprogramming as a mode of cancer drug resistance. Nature 546, 431–435 (2017).

10. Emert, B. L. et al. Variability within rare cell states enables multiple paths toward drug resistance. Nat. Biotechnol. 39, 865–876 (2021).

11. Yang, C., Tian, C., Hoffman, T. E., Jacobsen, N. K. & Spencer, S. L. Melanoma subpopulations that rapidly escape MAPK pathway inhibition incur DNA damage and rely on stress signalling. Nat. Commun. 12, 1747 (2021).

12. Elgin, S. C. R. & Reuter, G. Position-effect variegation, heterochromatin formation, and gene silencing in Drosophila. Cold Spring Harb. Perspect. Biol. 5, a017780 (2013).

13. Wu, S. et al. Independent regulation of gene expression level and noise by histone modifications. PLoS Comput. Biol. 13, e1005585 (2017).

14. Weinberger, L. et al. Expression noise and acetylation profiles distinguish HDAC functions. Mol. Cell 47, 193–202 (2012).

15. Walters, M. C. et al. Enhancers increase the probability but not the level of gene expression. Proceedings of the National Academy of Sciences 92, 7125–7129 (1995).

16. Dar, R. D. et al. Transcriptional burst frequency and burst size are equally modulated across the human genome. Proc. Natl. Acad. Sci. U. S. A. 109, 17454–17459 (2012).

17. Larson, D. R. et al. Direct observation of frequency modulated transcription in single cells using light activation. Elife 2, e00750 (2013).

18. Senecal, A. et al. Transcription factors modulate c-Fos transcriptional bursts. Cell Rep. 8, 75–83 (2014).

19. Faure, A. J., Schmiedel, J. M. & Lehner, B. Systematic Analysis of the Determinants of Gene Expression Noise in Embryonic Stem Cells. Cell Systems 5, 471–484.e4 (2017).

20. Karlić, R., Chung, H.-R., Lasserre, J., Vlahovicek, K. & Vingron, M. Histone modification levels are predictive for gene expression. Proc. Natl. Acad. Sci. U. S. A. 107, 2926–2931 (2010).

21. Akhtar, W. et al. Chromatin position effects assayed by thousands of reporters integrated in parallel. Cell 154, 914–927 (2013).

22. Kundaje, A. et al. Integrative analysis of 111 reference human epigenomes. Nature 518, 317–330 (2015).

23. Dey, S. S., Foley, J. E., Limsirichai, P., Schaffer, D. V. & Arkin, A. P. Orthogonal control of expression mean and variance by epigenetic features at different genomic loci. Mol. Syst. Biol. 11, 806 (2015).

24. Zhang, T., Foreman, R. & Wollman, R. Identifying chromatin features that regulate gene expression distribution. Sci. Rep. 10, 20566 (2020).

25. Elowitz, M. B., Levine, A. J., Siggia, E. D. & Swain, P. S. Stochastic gene expression in a single cell. Science 297, 1183–1186 (2002).

26. Ozbudak, E. M., Thattai, M., Kurtser, I., Grossman, A. D. & van Oudenaarden, A. Regulation of noise in the expression of a single gene. Nat. Genet. 31, 69–73 (2002).

27. das Neves, R. P. et al. Connecting variability in global transcription rate to mitochondrial variability. PLoS Biol. 8, e1000560 (2010).

28. Stewart-Ornstein, J., Weissman, J. S. & El-Samad, H. Cellular noise regulons underlie fluctuations in Saccharomyces cerevisiae. Mol. Cell 45, 483–493 (2012).

29. Sanchez, A. & Golding, I. Genetic determinants and cellular constraints in noisy gene expression. Science 342, 1188–1193 (2013).

30. Raser, J. M. & O‘Shea, E. K. Noise in gene expression: origins, consequences, and control. Science 309, 2010–2013 (2005).

31. Zopf, C. J., Quinn, K., Zeidman, J. & Maheshri, N. Cell-cycle dependence of transcription dominates noise in gene expression. PLoS Comput. Biol. 9, e1003161 (2013).

32. Hoffman, M. M. et al. Integrative annotation of chromatin elements from ENCODE data. Nucleic Acids Res. 41, 827–841 (2013).

33. Vallania, F. L. M. et al. Origin and consequences of the relationship between protein mean and variance. PLoS One 9, e102202 (2014).

34. Bar-Even, A. et al. Noise in protein expression scales with natural protein abundance. Nat. Genet. 38, 636–643 (2006).

35. ENCODE Project Consortium. An integrated encyclopedia of DNA elements in the human genome. Nature 489, 57–74 (2012).

36. Fu, A. Q. & Pachter, L. Estimating intrinsic and extrinsic noise from single-cell gene expression measurements. Stat. Appl. Genet. Mol. Biol. 15, 447–471 (2016).

37. Litzenburger, U. M. et al. Single-cell epigenomic variability reveals functional cancer heterogeneity. Genome Biol. 18, 15 (2017).

38. Moudgil, A. et al. Self-Reporting Transposons Enable Simultaneous Readout of Gene Expression and Transcription Factor Binding in Single Cells. Cell 182, 992–1008.e21 (2020).

39. Aznauryan, E. et al. Discovery and validation of human genomic safe harbor sites for gene and cell therapies. Cell Reports Methods vol. 2 100154 (2022).

40. Badia-i-Mompel, P. et al. decoupleR: ensemble of computational methods to infer biological activities from omics data. Bioinformatics Advances 2, vbac016 (2022).

41. Papapetrou, E. P. & Schambach, A. Gene Insertion Into Genomic Safe Harbors for Human Gene Therapy. Mol. Ther. 24, 678–684 (2016).

42. Bonny, A. R., Fonseca, J. P., Park, J. E. & El-Samad, H. Orthogonal control of mean and variability of endogenous genes in a human cell line. Nat. Commun. 12, 292 (2021).

43. Raj, A., Peskin, C. S., Tranchina, D., Vargas, D. Y. & Tyagi, S. Stochastic mRNA synthesis in mammalian cells. PLoS Biol. 4, e309 (2006).

44. Benzinger, D. & Khammash, M. Pulsatile inputs achieve tunable attenuation of gene expression variability and graded multi-gene regulation. Nat. Commun. 9, 3521 (2018).

45. Michaels, Y. S. et al. Precise tuning of gene expression levels in mammalian cells. Nat. Commun. 10, 818 (2019).

46. Pavani, G. & Amendola, M. Targeted Gene Delivery: Where to Land. Front Genome Ed 2, 609650 (2020).

47. Cao, J. et al. Comprehensive single-cell transcriptional profiling of a multicellular organism. Science 357, 661–667 (2017).

48. Rosenberg, A. B. et al. Single-cell profiling of the developing mouse brain and spinal cord with split-pool barcoding. Science 360, 176–182 (2018).

49. Qi, Z. et al. An optimized, broadly applicable piggyBac transposon induction system. Nucleic Acids Res. 45, e55 (2017).

50. Li, H. & Durbin, R. Fast and accurate short read alignment with Burrows-Wheeler transform. Bioinformatics 25, 1754–1760 (2009).

51. Rouhanifard, S. H. et al. ClampFISH detects individual nucleic acid molecules using click chemistry-based amplification. Nat. Biotechnol. (2018) doi:10.1038/nbt.4286.

52. Lawrence, M. et al. Software for computing and annotating genomic ranges. PLoS Comput. Biol. 9, e1003118 (2013).

53. Gu, Z., Eils, R., Schlesner, M. & Ishaque, N. EnrichedHeatmap: an R/Bioconductor package for comprehensive visualization of genomic signal associations. BMC Genomics 19, 234 (2018).

54. Bailey, T. L. STREME: Accurate and versatile sequence motif discovery. Bioinformatics (2021) doi:10.1093/bioinformatics/btab203.

55. Gupta, S., Stamatoyannopoulos, J. A., Bailey, T. L. & Noble, W. S. Quantifying similarity between motifs. Genome Biol. 8, R24 (2007).

56. Durand, N. C. et al. Juicer Provides a One-Click System for Analyzing Loop-Resolution Hi-C Experiments. cels 3, 95–98 (2016).

57. Bianchi, V. et al. Detailed Regulatory Interaction Map of the Human Heart Facilitates Gene Discovery for Cardiovascular Disease. bioRxiv 705715 (2019) doi:10.1101/705715.

58. Quinlan, A. R. & Hall, I. M. BEDTools: a flexible suite of utilities for comparing genomic features. Bioinformatics 26, 841–842 (2010).

59. Grant, C. E., Bailey, T. L. & Noble, W. S. FIMO: scanning for occurrences of a given motif. Bioinformatics 27, 1017–1018 (2011).

60. Harmston, N., Ing-Simmons, E., Perry, M., Barešić, A. & Lenhard, B. GenomicInteractions: An R/Bioconductor package for manipulating and investigating chromatin interaction data. BMC Genomics 16, 963 (2015).

61. Satopaa, V., Albrecht, J., Irwin, D. & Raghavan, B. Finding a ‘Kneedle’ in a Haystack: Detecting Knee Points in System Behavior. in 2011 31st International Conference on Distributed Computing Systems Workshops 166–171 (2011).

62. Wolf, F. A., Angerer, P. & Theis, F. J. SCANPY: large-scale single-cell gene expression data analysis. Genome Biol. 19, 15 (2018).

